# A Python Dash App and cPanel workflow to automate metabolomics data analyses and visualisation

**DOI:** 10.64898/2026.05.01.722139

**Authors:** Jessica M. O’Loughlin, Tessa Moses

## Abstract

Metabolomics offers a sophisticated analytical framework for characterising the molecular phenotype of biological organisms and complex living systems at a high resolution. As the functional endpoint of the omics cascade, the metabolome serves as a close reflection of cellular activity. It integrates genetic, transcriptomic and proteomic variations with external environmental influences. However, the inherent complexity of metabolomic datasets, characterised by high-dimensional chemical diversity, wide dynamic ranges, and significant matrix effects, necessitates a rigorous suite of chemometric and bioinformatic workflows. For researchers uninitiated in computational biology, the multi-stage requirement for raw data pre-processing, signal deconvolution, and multivariate statistical modelling (such as PCA or PLS-DA) presents a substantial barrier to entry. Navigating these convoluted data architectures remains a primary challenge in deriving biological meaning from the global metabolic profile. Here, we present a workflow to use Python Dash Apps to create a user-friendly interface for simplifying data processing and statistical calculations. Users can select their desired samples to initiate calculations for various statistical tests, generating interactive and publication-quality figures to explore their results. These apps were deployed on an Apache server via cPanel, allowing individuals to share their findings with collaborators and for research facilities to share metabolomics results with their users.

## Introduction

Metabolomics coupled with high-resolution mass spectrometry (MS) has emerged as an exceptionally versatile and high-throughput analytical approach. It allows for the in-depth exploration of molecular phenotypes across a vast spectrum of biological systems and diverse organismal models (Doerr, 2017; Dörr et al., 2013; Fessenden, 2016). As the functional downstream endpoint of the central dogma, the metabolome provides a proximal and real-time reflection of cellular physiology (Hautbergue et al., 2018; Ramirez-Gaona et al., 2017; Sajed et al., 2016; Wishart et al., 2018). By identifying and semi-quantifying thousands of discrete metabolites, ranging from polar to semi-polar primary metabolites and complex non-polar lipids, this approach yields invaluable insights into the metabolic flux and molecular dynamics that underpin critical environmental (Brown et al., 2024; Swenson et al., 2018; Thukral et al., 2023; Yongliang et al., 2022), clinical (Chakaroun et al., 2026; Flores et al., 2024; Han et al., 2021; Mayo-Martínez et al., 2021; Suri et al., 2023), and industrial (Cortada-Garcia et al., 2024; Safo et al., 2021; Tanaka et al., 2024; Yu et al., 2022) bioprocesses.

Despite the descriptive power of metabolomics, the transition from raw spectral acquisition to biological interpretation is fraught with significant computational challenges. The resulting datasets are inherently high-dimensional and susceptible to various forms of analytical noise, including matrix effects, signal drift over long injection sequences, and variations in sample concentration. To mitigate these artifacts, data must undergo rigorous pre-processing steps, including peak alignment, deconvolution, and sophisticated normalisation strategies (De Livera et al., 2012; Li et al., 2017; Sun & Xia, 2024). These steps are foundational; without appropriate scaling and transformation, the biological signal can easily be obscured by technical variance, leading to erroneous statistical conclusions during subsequent univariate or multivariate visualisations.

While several sophisticated software suites (Schmid et al., 2023; Smith et al., 2006), R-based packages (Idkowiak et al., 2025; Liang et al., 2025; Mock et al., 2018; Nicolotti et al., 2021), and web-based NetWare tool (Pang et al., 2024) has been developed to facilitate these workflows, a significant barrier to entry remains. In our experience, the steep learning curve associated with command-line interfaces or complex parameter tuning can be daunting for researchers whose primary expertise lies in biology or clinical medicine rather than bioinformatics. This technical friction often slows the pace of discovery and limits the accessibility of metabolomics to specialised laboratories. To address this gap, we present a user-centric interactive report designed to democratise the metabolomics pipeline. Our tool simplifies the most taxing elements of the process, specifically data normalisation and the generation of complex statistical architectures, into a seamless, intuitive interface. By abstracting the underlying code, the tool allows users to focus on the biological relevance of their data; researchers need only select the specific sample cohorts they wish to compare, and the system automatically executes the necessary statistical calculations and high-fidelity visualisations. This approach ensures that robust, reproducible metabolomic analysis is accessible to the broader scientific community, regardless of their prior computational experience.

The interactive reports are scripted using Python, a high-level, interpreted, and object-oriented programming language that has become a lingua franca of the global scientific community. Python’s dominance in bioinformatics is driven by its expansive ecosystem of specialised libraries, which provide the robust data structures required to standardise the complex mathematical operations inherent in metabolomics.

These operations, ranging from missing value imputation and probabilistic quotient normalisation to multivariate projections, are essential for transforming raw spectral data into interpretable biological signals (Pedregosa et al., 2018; Virtanen et al., 2020). Furthermore, Python’s integration with advanced visualisation libraries allows for the generation of publication-quality figures that remain fully interactive, enabling researchers to interrogate individual data points within a larger global profile (Nicolas Kruchten; Andrew Seier; Chris Parmer, 2025).

To bridge the gap between back-end computation and front-end usability, we utilised the Dash framework by Plotly. Dash applications are reactive web environments programmed entirely in Python, eliminating the need for separate JavaScript or HTML development while providing a sophisticated interface for data interaction. These Dash Apps are uniquely suited for metabolomics because they allow for real-time filtering, hover-over metadata display, and dynamic scaling of statistical plots. By hosting these applications on a publicly accessible server, the resulting reports transcend the limitations of traditional static files. Collaborators can explore the data synchronously from any location, removing the logistical burden of emailing large, disparate PDFs or Excel spreadsheets and ensuring a single source for the dataset.

While the technical advantages of Dash are clear, the deployment phase often presents a significant financial and administrative hurdle. Currently, the industry-standard recommendations for hosting these applications include Plotly Cloud (*Plotly Cloud*, 2026) or Dash Enterprise (*Dash Enterprise*, 2026) with Plotly Dash (*Publishing Dash Apps*, 2026). While these platforms offer high performance and ease of use, they typically operate under a software-as-a-service (SaaS) model requiring recurring subscription fees. For many academic laboratories, individual researchers, or small-scale industrial projects, these costs can be prohibitive, effectively limiting the accessibility and long-term sustainability of interactive data sharing.

To address these accessibility constraints, we have identified a more cost-effective pathway using cPanel, a widely utilised web hosting control panel prevalent in many institutional and commercial web hosting environments. Although cPanel is traditionally associated with static website hosting, it has robust, underutilised functionalities for supporting persistent Python applications via the WSGI (Web Server Gateway Interface). Therefore, in addition to the interactive reporting tool detailed in this article, we provide a comprehensive methodology for deploying these Dash applications via the cPanel interface. This approach provides a scalable, low-cost alternative to enterprise-level servers, empowering researchers to host and share their interactive metabolomic findings using existing web infrastructure.

## Methodology

### Software architecture and environment requirements

The interactive reporting tool was architected using Python (v3.9), leveraging the Dash (v2.17.1) framework by Plotly for reactive web development. To ensure reproducibility and facilitate deployment across diverse computing environments, a comprehensive list of library dependencies is maintained in the project’s associated GitHub repository (Jessica O’Loughlin; Tessa Moses, 2026)

### Data structure and compatibility

To maintain interoperability with existing metabolomics bioinformatic pipelines, the application was engineered to support the standardised comma-separated values (CSV) format. The input architecture requires an ‘unpaired samples in columns’ configuration, featuring peak intensities and metadata annotations consistent with the requirements for MetaboAnalyst 6.0 (Pang et al., 2024). This design choice ensures that researchers can transition seamlessly from raw data processing to advanced visualisation without the need for additional file reformatting.

### Access modalities and deployment strategies

The application offers two distinct functional modalities for data ingestion, designed to accommodate different collaborative and analytical workflows. In the static directory mapping (server-side loading) configuration, the dataset path is defined globally within the application script. This pre-loaded approach is optimised for deployment scenarios where a specific, curated dataset needs to be shared securely with collaborators via a hosted URL (Figure 1a). The second workflow uses dynamic user upload (client-side ingestion). This modality utilises the dcc.Upload component (Nicolas Kruchten; Andrew Seier; Chris Parmer, 2025), providing a drag-and-drop interface for end-users to upload local CSV files directly into the browser session. This flexible architecture is better suited for general-purpose analytical environments where the same application instance is used to interrogate multiple, independent metabolomics datasets (Figure 1b).

**Figure 1.**
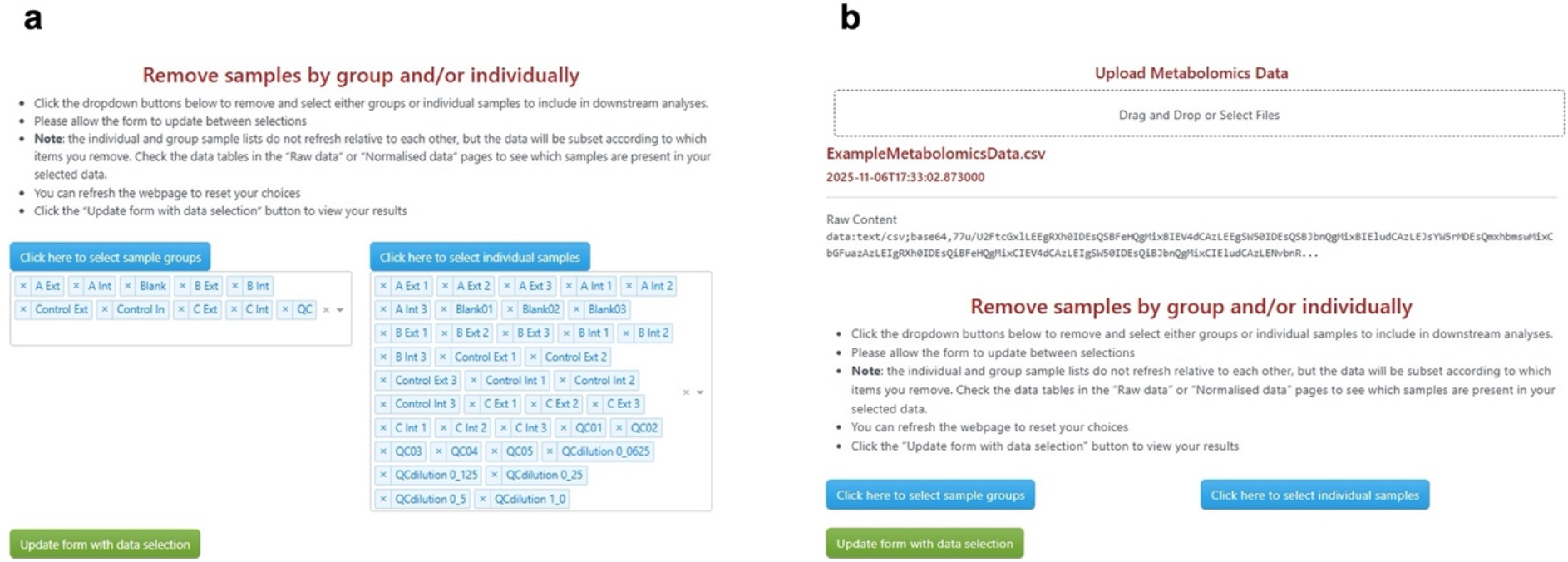
The two distinct access modalities used for data deployment. (**a**) Data is pre-loaded by specifying the file directory within the script. (**b**) Data is uploaded by the user to the interface.

### Data pre-processing and normalisation

Raw peak intensity metabolomics data were subjected to a rigorous normalisation pipeline to achieve a Gaussian distribution. This process included a Log10 transformation, mean-centring, and Pareto scaling (Sun & Xia, 2024). Users can evaluate the data distribution and absolute values both pre- and post-normalisation in the ‘Raw Data’ and ‘Normalised Data’ tabs within the reporting interface. Comprehensive processing scripts are available via our GitHub repository (Jessica O’Loughlin; Tessa Moses, 2026).

### Statistical analysis and multivariate modelling

Multivariate statistical analyses were performed using the Scikit-learn module (Pedregosa et al., 2018). Data reduction was achieved through principal component analysis (PCA), which was utilised as an unsupervised technique to identify inherent groupings and outliers, and partial least squares – discriminant analysis (PLS-DA), which was employed as a supervised method to maximise class separation. Additionally, Scikit-learn was used for data scaling during the generation of heat map visualisations. Univariate analysis, specifically the calculation of p-values for volcano plots, was conducted using the t-test function from the SciPy module (Virtanen et al., 2020). The accuracy of these computational outputs was validated against results generated via MetaboAnalyst (Pang et al., 2024), ensuring consistent analytical performance.

### Software deployment and Dash application

Interactive visualisations were deployed as a Python Dash application using the cPanel interface (*CPanel, WebHost Manager*, 2026), within the Setup Python App section. To ensure successful deployment, the automatically generated passenger_wsgi.py file was manually modified to bridge the Dash server with the web interface. Detailed parameters, initialisation steps, and the modified WSGI configurations are documented in the associated GitHub repository (Jessica O’Loughlin; Tessa Moses, 2026).

## Results and Discussion

### Automated statistical workflows and normalisation validation

Upon submission of the selected sample cohorts, the application executes the pre-defined statistical pipeline, generating high-fidelity graphical outputs. A critical component of this workflow is the normalisation validation interface (Figure 2). By providing a side-by-side comparison of raw versus processed peak intensities, the platform allows users to verify that the Log10 transformation and Pareto scaling have successfully corrected for heteroscedasticity and achieved a Gaussian distribution. This transparency is vital for establishing the statistical validity of downstream multivariate analyses, ensuring that biological variation is not obscured by technical noise.

**Figure 2.**
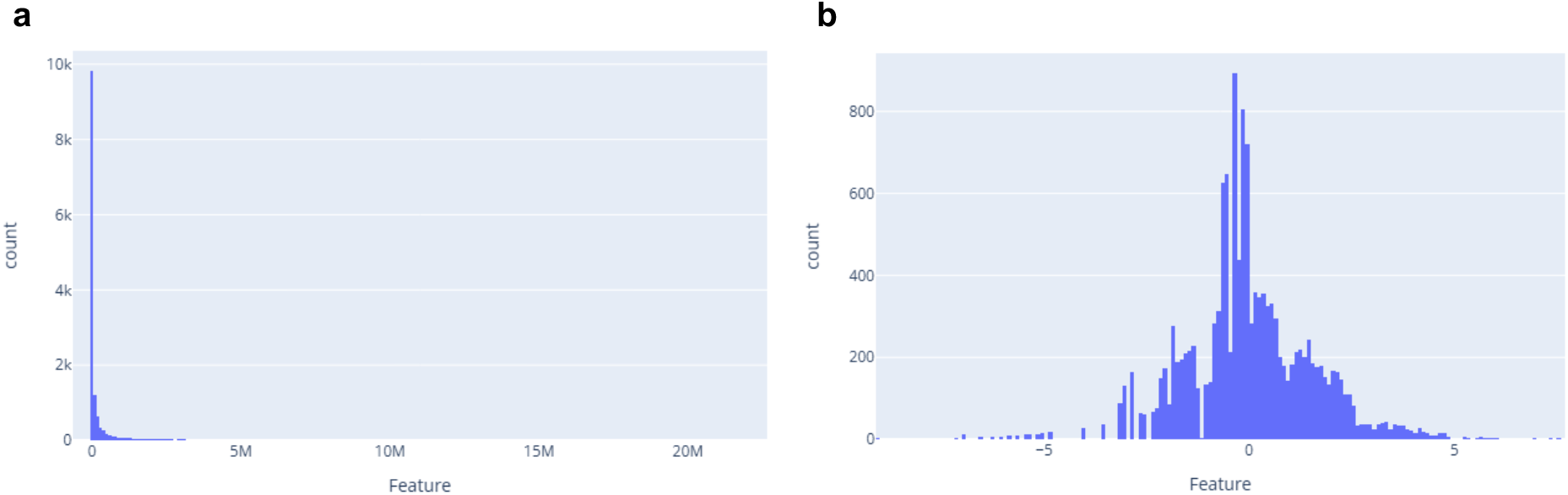
Example of the distribution of feature intensities (**a**) in the raw data and (**b**) after it is normalised by Log_10_ transformation and Pareto scaling in the same dataset.

### Enhanced interactivity and multi-dimensional data mining

Unlike static reporting tools, the integration of Plotly-based web elements (Nicolas Kruchten; Andrew Seier; Chris Parmer, 2025) enables a deep dive into the metabolomics dataset. Every data point within the PCA and PLS-DA scores and loadings plots is interactive (Figure 3a-b). Hovering over specific coordinates reveals metadata and sample-specific identifiers, allowing for the rapid identification of biological outliers or transitionary phenotypes. Additionally, selecting a data point in the loadings plot enables the user to evaluate the intensities of that metabolite across the previously selected sample groups (Figure 3c).

**Figure 3.**
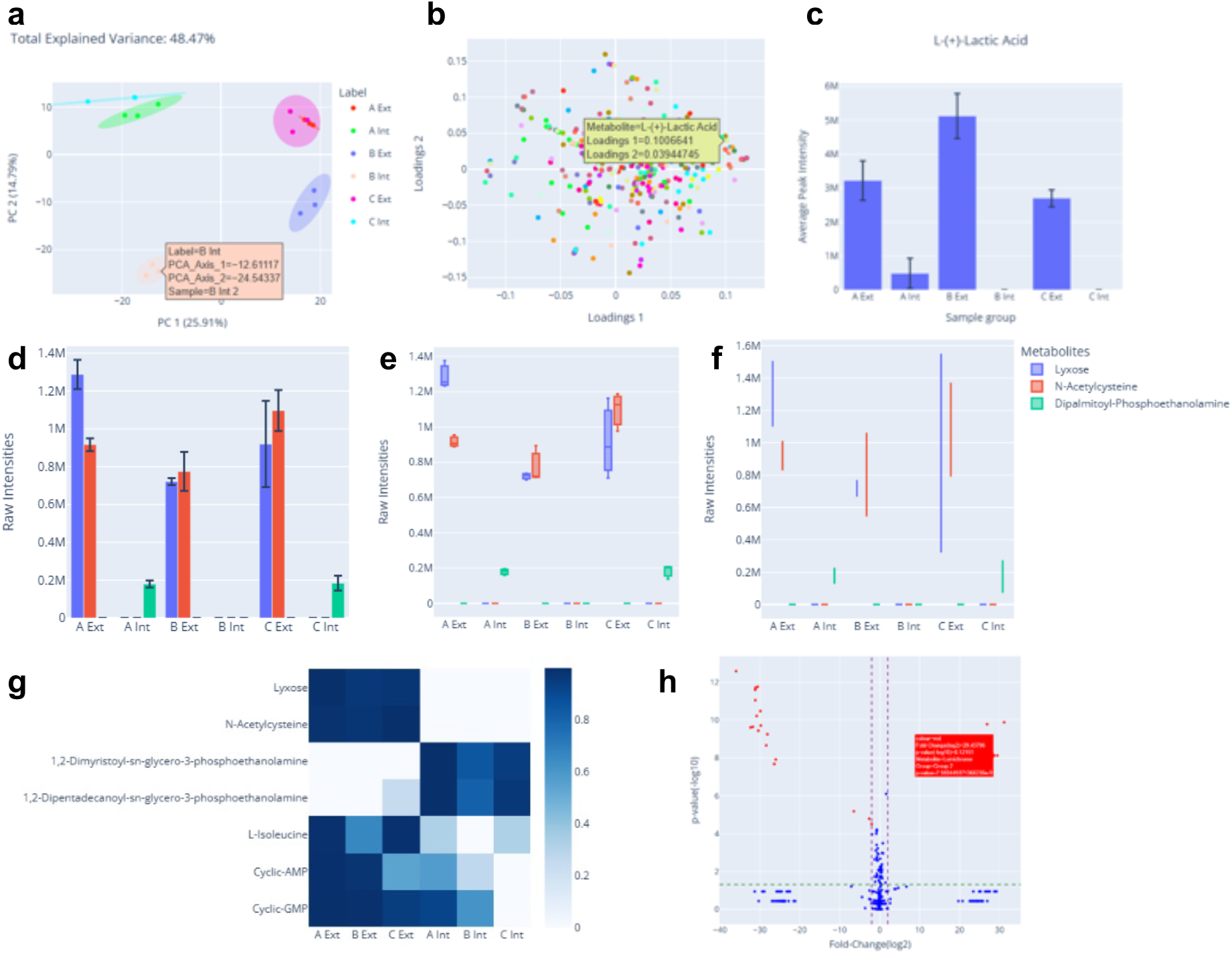
The interactive multi-dimensional data mining tools. (**a**) The PCA 2D scores plot shows separation of the experimental groups. (**b**) Loadings plot associated with the PCA separation and (**c**) the bar chart for the selected data point, L-(+)-lactic acid. (**d**) Bar chart, (**e**) box and whisker plot and (**f**) violin plot for lyxose, N-acetylcysteine and dipalmitoyl phosphoethanolamine. (**g**) Heatmap of selected metabolites. (**h**) Volcano plot showing log2 fold change on X-axis and −log10 p-value on Y-axis. The user-selected significance thresholds are shown with the horizontal and vertical dashed lines.

To further facilitate data communication, we implemented custom-coded functions designed for comparative analysis. Multi-metabolite comparisons allow users to simultaneously plot the relative abundances of multiple metabolites across different experimental groups. This feature allows identifying and visualising co-regulated pathway components. The data can be visualised as linear or Log10-transformed raw or normalised intensities in these plots. In addition, the user has a choice of visualising the data as (i) bar charts for comparing quantities between groups (Figure 3d), (ii) box and whisker plots (Figure 3e), which represent the interquartile range (IQR) and the middle 50% of the data, or (iii) violin plots (Figure 3f), which show the probability density of the data at different values.

Heatmaps offer an intuitive, colour-coded method to identify patterns, trends, and anomalies within large, complex datasets such as metabolomics. The dynamic heatmap generation tool is a customisable heatmap module (Figure 3g) that employs Scikit-learn scaling to visualise global metabolic shifts, allowing users to cluster samples based on shared chemical signatures.

Volcano plots are a powerful visualisation tool commonly used in metabolomics experiments to identify significant changes between two experimental conditions. They combine a measure of statistical significance (p-value) with the magnitude of change (fold change) in a single scatter plot. The responsive volcano plot interface features adjustable significance and fold-change thresholds, which are user-defined. Data points are colour-coded responsively (Figure 3h), enabling immediate visual confirmation of the most statistically significant metabolic perturbations.

### Data portability and collaborative utility

A significant hurdle in metabolomics is the siloing of data. To address this, we integrated a direct-to-Excel export function for every generated visualisation. This ensures that the underlying processed values remain portable, allowing researchers to re-format figures for specific publication requirements or integrate them into proprietary software.

When deployed via the cPanel/WSGI framework, the application serves as a centralised resource. It democratises data access by allowing multiple stakeholders, from lab technicians to principal investigators, to interact with the same processed dataset independently. This architecture eliminates the common discrepancy issues found when different researchers apply varied parameters to the same raw data, thereby ensuring absolute consistency in both processing and biological interpretation across the research team.

## Data availability

The code and an example dataset for the Python Dash App and the functions used to perform the calculations can be found in our GitHub repository https://github.com/EdinOmics/A-Python-Dash-App-and-cPanel-workflow-to-automate-metabolomics-data-analyses-and-visualisation.

## Acknowledgements

The authors thank Alain Forrester and Colin McLaren from the University of Edinburgh for their invaluable support in troubleshooting early deployment issues.

